# Impact of b-value on estimates of apparent fibre density

**DOI:** 10.1101/2020.01.15.905802

**Authors:** Sila Genc, Chantal M.W. Tax, Erika P. Raven, Maxime Chamberland, Greg D. Parker, Derek K. Jones

## Abstract

Recent advances in diffusion magnetic resonance imaging (dMRI) analysis techniques have improved our understanding of fibre-specific variations in white matter microstructure. Increasingly, studies are adopting multi-shell dMRI acquisitions to improve the robustness of dMRI-based inferences. However, the impact of b-value choice on the estimation of dMRI measures such as apparent fibre density (AFD) derived from spherical deconvolution is not known. Here, we investigate the impact of b-value sampling scheme on estimates of AFD. First, we performed simulations to assess the correspondence between AFD and simulated intra-axonal signal fraction across multiple b-value sampling schemes. We then studied the impact of sampling scheme on the relationship between AFD and age in a developmental population (n=78) aged 8-18 (mean=12.4, SD=2.9 years) using hierarchical clustering and whole brain fixel-based analyses. Multi-shell dMRI data were collected at 3.0T using ultra-strong gradients (300 mT/m), using 6 diffusion-weighted shells ranging from 0 – 6000 s/mm^2^. Simulations revealed that the correspondence between estimated AFD and simulated intra-axonal signal fraction was improved with high b-value shells due to increased suppression of the extra-axonal signal. These results were supported by *in vivo* data, as sensitivity to developmental age-relationships was improved with increasing b-value (b=6000 s/mm^2^, median R^2^ = .34; b=4000 s/mm^2^, median R^2^ = .29; b=2400 s/mm^2^, median R^2^ = .21; b=1200 s/mm^2^, median R^2^ = .17) in a tract-specific fashion. Overall, estimates of AFD and age-related microstructural development were better characterised at high diffusion-weightings due to improved correspondence with intra-axonal properties.

## 1. Introduction

Diffusion magnetic resonance imaging (dMRI; Le Bihan and Breton (1985)) offers a magnified window into white matter by probing the tissue microstructure properties. Various dMRI modelling and analysis techniques are available, which aim to summarise the local architecture of white matter as a quantitative metric. However, the biological interpretations around commonly investigated dMRI metrics rests heavily on whether the acquisition protocol can capture the relevant microstructural attributes (Lebel and Deoni, 2018; Tournier, et al., 2011).

Traditionally, studies have acquired dMRI data with one diffusion-weighting (or b-value shell), opting for either low b-values (e.g. b = 1000 s/mm^2^) for diffusion tensor imaging (DTI) analyses (Jones, et al., 1999; Landman, et al., 2007), or moderate-to-high b-values (e.g. b ≥ 3000 s/mm^2^) for probabilistic tractography (Tournier, et al., 2013). More recently, with the advent of multi-slice accelerated imaging (Barth, et al., 2016), the acquisition of multiple dMRI shells has become more feasible. This has considerably improved data acquisition capabilities for sensitive populations (such as children and clinical populations) which may not withstand long acquisition times (Kunz, et al., 2014; Silk, et al., 2016; Somerville, et al., 2018).

Multi-shell dMRI data has been used in conjunction with various analysis approaches (Novikov, et al., 2018; Zhang, et al., 2012) across a variety of applications (Genc, et al., 2018a; Kunz, et al., 2014; Pines, et al., 2019). Measures derived from constrained spherical deconvolution (CSD; Dell’Acqua and Tournier (2019)) can infer the intra-axonal signal fraction along multiple fibre pathways (Dell’Acqua, et al., 2013; Raffelt, et al., 2012). One such measure of microstructural organisation, termed apparent fibre density (AFD), can indicate relative differences in the white matter fibre density per unit volume of tissue. Given that the specificity to the intra-axonal water signal is maximised at high b-values due to higher restriction of water diffusion (Figure 1), AFD can be sensitive to axon density at high diffusion-weightings (Raffelt, et al., 2012).

**Figure 1:**
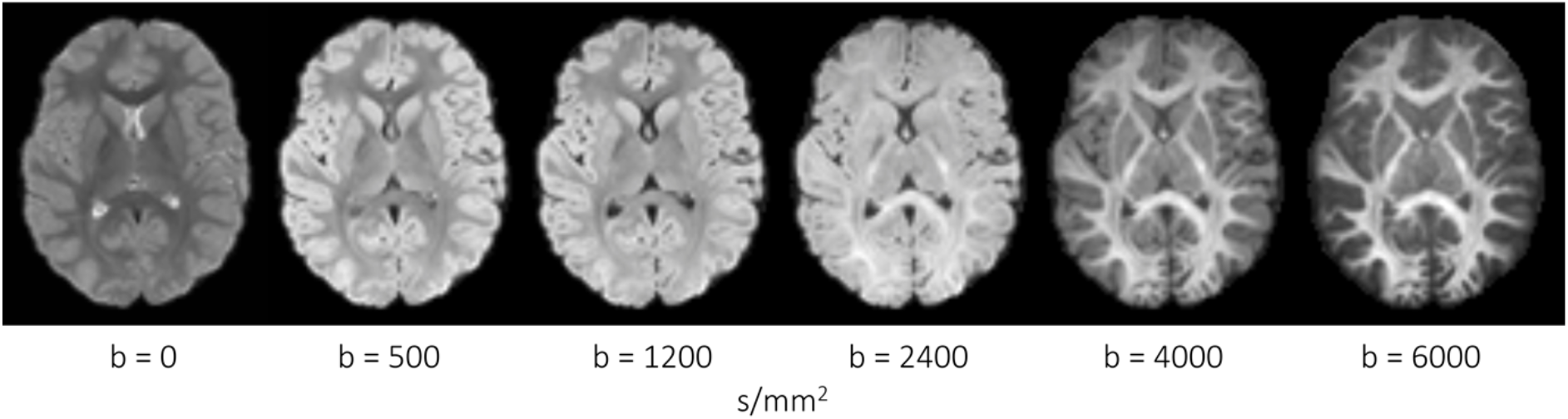
Spherical harmonics (zero order) maps derived from a representative participant (age 8 years). Visually, increasing b-value from 0 – 6000 s/mm^2^ leads to greater specificity to the signal attributed to the intra-axonal space.

Analysis frameworks such as fixel-based analysis (FBA; Raffelt, et al. (2017)) provide a means to test fibre-specific differences in AFD within a population. FBA offers two major advantages over alternative dMRI analysis techniques: sensitivity to fibre properties (density and morphology), and specificity to *fibre* populations within *voxels* (or *‘fixels’*). This combination of improved sensitivity and specificity increases the possibility of assigning group differences in fibre properties to specific fibre populations (Dimond, et al., 2019; Gajamange, et al., 2018; Genc, et al., 2018b; Mito, et al., 2018).

In practise, FBA is compatible with both single-shell (Dhollander, et al., 2016) and multi-shell (Jeurissen, et al., 2014) dMRI data. An intuitive choice might be to use all available dMRI data to compute fibre-specific AFD. However, this might not be compatible with the underlying assumptions of AFD reflecting intra-axonal properties. In addition, sensitivity to the extra-axonal signal upon the inclusion of lower b-values can influence the response function choice, resulting in a potential mismatch between the response function and the true underlying fibre properties.

Combining FBA with the very latest in MRI gradient hardware (300 mT/m) (Jones, et al., 2018), we explore the impact of sampling scheme on AFD estimates using a rich developmental dataset comprising multi-shell diffusion MRI data with b-values ranging from 0 – 6000 s/mm^2^. Firstly, we simulate multiple fibre geometries to showcase how discrepancies in ‘true’ microstructural configurations can influence the interpretations of AFD generated from both single-shell and multishell dMRI data. We then conduct experiments to confirm the theory that AFD is more sensitive and specific to axon density at higher b-values, demonstrated by sensitivity to detecting age-relationships in a developmental population of children and adolescents.

## 2. Methods

### 2.1. Simulations

Single fibre populations were simulated with the intra- and extra-axonal spaces represented by axially symmetric tensors; the second and third eigenvalues were set to zero for the intra-axonal tensor and equal but non-zero for the extra-axonal tensor (Jespersen, et al., 2007; Kroenke, et al., 2004). The intra-axonal and extra-axonal parallel diffusivities were set to 1.9 *μm*^2^/*ms*, and 42 different combinations were simulated with intra-axonal signal fraction *f* = [0.2,0.3,0.4,0.5,0.6,0.7,0.8] and extra-axonal perpendicular diffusivity *D*_*e*,⊥_ = [0.2,0.4,0.6,0.8,1,1.2] *μm*^2^/*ms*. 100 Rician noise generalisations were computed with three different signal-to-noise ratio (SNR) values on the b=0 signal (SNR = 50; 35; and 20). The response function, which should reflect the properties of a single fibre population (Tax, et al., 2014), was set to have *f* = 0.3 and *D*_*e*,⊥_ = 0.8 *μm*^2^/*ms* informed by values estimated from the group-wise response function used in this study. These values are in the range of previously reported estimates of white matter *in vivo* (Fieremans, et al., 2011; Novikov, et al., 2018).

### 2.2. Participants

We scanned a sample of typically developing children aged 8-18 years recruited as part of the Cardiff University Brain Research Imaging Centre (CUBRIC) Kids study (Genc, et al., 2019; Raven, et al., 2019). This study was approved by the School of Psychology ethics committee at Cardiff University. Participants and their parents/guardians were recruited via public outreach events. Written informed consent was provided by the primary caregiver of each child participating in the study, and adolescents aged 16-18 years additionally provided written consent. Children were excluded from the study if they had non-removable metal implants, and if they reported history of a major head injury or epilepsy. All procedures were completed in accordance with the Declaration of Helsinki.

A total of 78 children between the ages of 8 – 18 years (Mean = 12.4, SD = 2.9 years) were included in the current study (45 female).

### 2.3. Diffusion magnetic resonance imaging

#### Image acquisition and pre-processing

Diffusion Magnetic Resonance Imaging (dMRI) data were acquired on a 3.0T Siemens Connectom system with ultra-strong (300 mT/m) gradients. Multi-shell dMRI data were collected using the following parameters: TE/TR = 59/3000 ms; voxel size = 2×2×2 mm; b-values = 0 (14 volumes, interleaved), 500 (30 directions), 1200 (30 directions), 2400 (60 directions), 4000 (60 directions), and 6000 (60 directions) s/mm^2^. The larger number of volumes across the higher diffusion weightings were to compensate for lower SNR and to capture the higher angular resolution present at higher b-values (Tournier, et al., 2013). Diffusion MRI data were acquired using electrostatic repulsion generalised across multiple shells (Caruyer, et al., 2013). Data were acquired in an anterior-posterior (AP) phase-encoding direction, with one additional PA volume. The total acquisition time (across four acquisition blocks) was 16 minutes and 14 seconds.

Pre-processing of dMRI data involved steps largely in line with recommended steps for standard 3.0T systems, interfacing various tools such as FSL (Smith, et al., 2004), MRtrix3 (Tournier, et al., 2019), and ANTS (Avants, et al., 2011). These steps included: denoising (Veraart, et al., 2016), slicewise outlier detection (SOLID; Sairanen, et al. (2018)), and correction for drift (Vos, et al., 2017); motion, eddy, and susceptibility-induced distortions (Andersson, et al., 2003; Andersson and Sotiropoulos, 2016); Gibbs ringing artefact (Kellner, et al., 2016); bias field (Tustison, et al., 2010); and gradient non-linearities (Glasser, et al., 2013; Rudrapatna, et al., 2018). Root mean squared (RMS) displacement from *eddy* (Andersson and Sotiropoulos, 2016) was used as a summary measure of global head motion. Estimates of SNR were performed by taking the signal in the white matter and dividing this by the signal outside of the brain (for each b=0 image). SNR estimates in the *in vivo* data were: mean = 48.02, SD = 7.46.

#### Image processing and analysis

To compare multiple sampling schemes, pre-processed dMRI data were further processed and analysed separately for each sampling scheme in a common population-template space, using a recommended framework (Raffelt, et al., 2017). Firstly, data were intensity normalised and spatially upsampled to 1.3 mm^3^ isotropic voxel size to increase anatomical contrast and improve tractography (Dyrby, et al., 2014). For single-shell (ss) single-tissue constrained spherical deconvolution (CSD), a fibre orientation distribution (FOD; Tournier, et al. (2007)) was estimated in each voxel with maximal spherical harmonics order *l_max_* = 8 for shells with high angular resolution (b=2400, 4000, 6000 – 60 directions each) and *l_max_* = 6 for shells with lower angular resolution (b=1200 – 30 directions). Multishell (ms) multi-tissue CSD was performed using a separate framework (Dhollander, et al., 2016; Jeurissen, et al., 2014). Following FOD estimation, we derived a population template using all diffusion volumes (msall), and subsequently registered subject-specific and sampling-scheme-specific FOD maps to this template (Figure S1). We then computed an apparent fibre density (AFD) map containing fibre-specific AFD along each fixel for each subject (Raffelt, et al., 2017).

In order to estimate AFD along various commonly investigated white matter fibre pathways, white matter tract segmentation was performed. We applied the automated *TractSeg* technique (Wasserthal, et al., 2018; Wasserthal, et al., 2019) in population template space, as this technique provides a balance between manual dissection and atlas-based tracking approaches. Of the existing library of 72 tracts, we chose to delineate 38 commonly investigated fibre pathways bilaterally for the left (L) and right (R) hemisphere (Figure S2). This included: AF: arcuate fasciculus; ATR: anterior thalamic radiation; CA: anterior commissure; CC: corpus callosum [1=rostrum, 2=genu, 3=rostral body, 4=anterior midbody, 5=posterior midbody; 6=isthmus, 7=splenium]; CG = cingulum; CST: corticospinal tract; FX: fornix; ICP: inferior cerebellar peduncle; IFOF: inferior fronto-occipital fasciculus; ILF: inferior longitudinal fasciculus; MCP: middle cerebellar peduncle; MLF: middle longitudinal fasciculus; OR: optic radiation; superior longitudinal fasciculus: SLF [I, II, III]; and UF: uncinate fasciculus. Each tractography map was converted to a fixel map to segment fixels corresponding to streamlines, and AFD was computed within each tract-specific fixel map for further statistical analysis.

### 2.4. Statistical analyses

#### Impact of b-value sampling scheme

Statistical analyses were performed within R (v3.4.3) and visualisations were carried out in RStudio (v1.2.1335). The coefficient of determination (R^2^) was computed to summarise the proportion of variance explained by age for each sampling-scheme in each tract. Linear models were computed, whereby AFD in each tract was entered as the dependent variable, age was entered as the independent variable, and sex and RMS displacement were set as nuisance variables. To compare sampling schemes in terms of their relationship with age, the difference in R^2^ was bootstrapped with 10,000 samples to compute 95% bias corrected accelerated (BCa) confidence intervals.

Hierarchical clustering was performed to discern clusters of sensitivity to age-relationships across various combinations of b-value sampling schemes and white matter tracts. These results were visualised as a heatmap with hierarchical clustering using the *‘gplots’* package (Warnes, et al., 2015) using Euclidean distance and complete agglomeration for clustering. To account for family-wise error (FWE) we made use of a strict Bonferroni correction by adjusting our p-value threshold by the 152 comparisons (38 tracts x 4 sampling schemes). As a result, our statistical significance was defined as *p* < 3.3e-4.

#### Whole-brain fixel-based analysis

Separate statistical analyses were performed for each single-shell sampling scheme (b = 1200; 2400; 4000; and 6000 s/mm^2^) using connectivity-based fixel enhancement (CFE), which provides a permutation-based, family-wise error (FWE) corrected p-value for every individual fixel in the template image (Raffelt, et al., 2015). For each sampling scheme, we tested the relationship between AFD and age, covarying for sex. For these whole-brain analyses, statistical significance was defined as *p_FWE_* < .05. Statistically significant fixels were converted into binary fixel maps, and an intersection mask was computed to quantify the proportion of overlapping fixels between sampling schemes.

## 3. Results

### 3.1. Simulations

The results of the simulations for AFD across various fibre geometries and sampling schemes is summarised in Figure 2. Compared to the highest single shell acquisition (ss_6000_), we observe a statistically significant three-way interaction between *D*_*e*,⊥_, *f*, and sampling-scheme for ss_1200_: β [95% CI] = .80 [.44, 1.2]; ss_2400_: β [95% CI] = .55 [.19, .91]; and ms_all_: β [95% CI] = .83 [.47, 1.2]. These observed differences are visually reflected by a greater dependency of AFD and *f* on simulated *D*_*e*,⊥_ as a result of the discrepancy with the response function.

**Figure 2:**
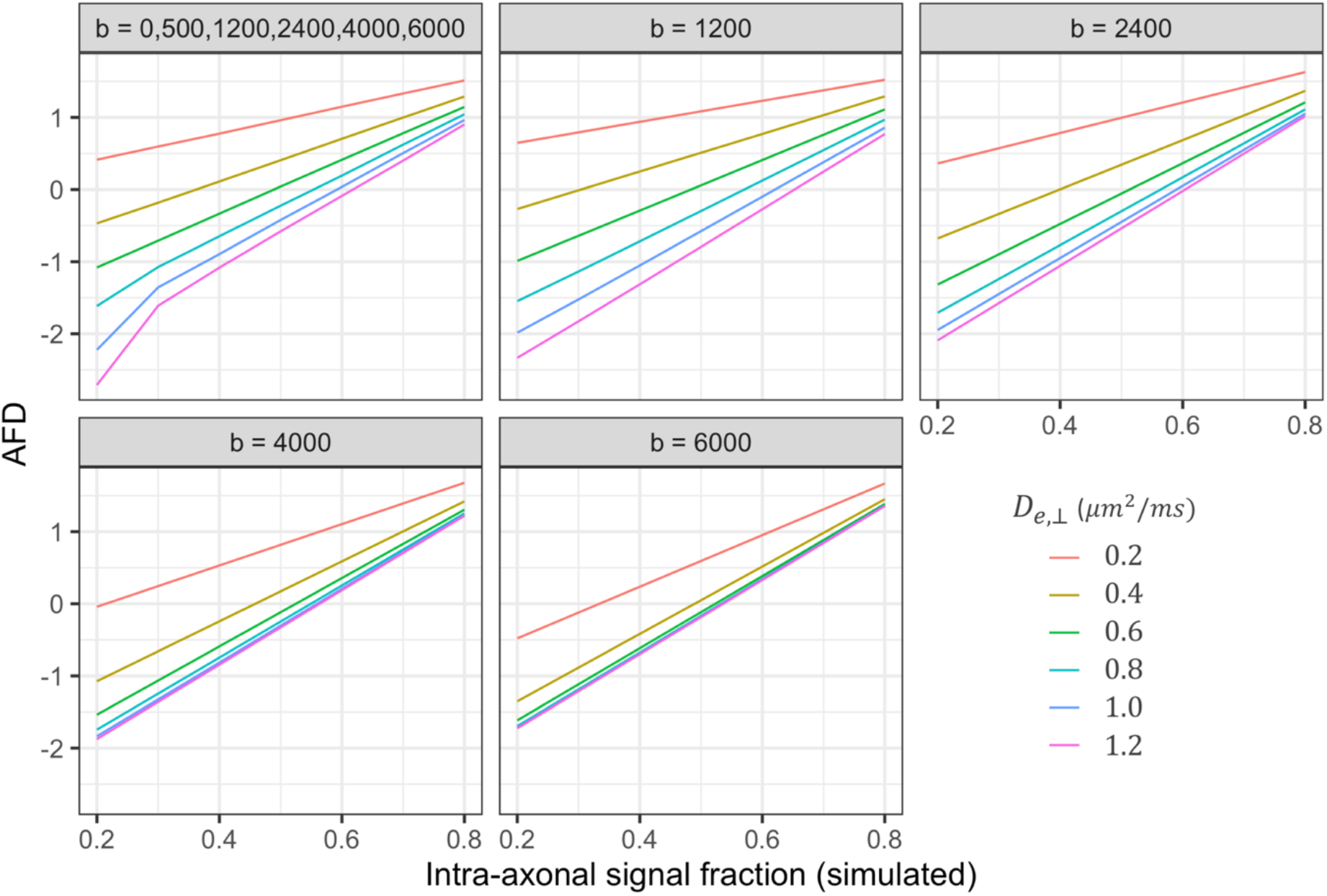
AFD for simulated fibre geometries across five sampling schemes (b-value, s/mm^2^). Variations to simulated intra-axonal signal fraction and perpendicular diffusivity of the extra-axonal space (*D*_*e*,⊥_) were tested to compare AFD across multiple fibre geometries.

When considering the full multi-shell acquisition (ms_all_) there are multiple degenerate scenarios whereby different combinations of *f* and *D*_*e*,⊥_ could result in the same AFD, compared with high b-value shells (i.e. ss_4000_ or ss_6000_). From the simulated scenarios for example, if AFD (ms_all_) = 1.2, it can be seen that there are at least six combinations of *f* and *D*_*e*,⊥_ resulting in this value (Figure 2). Whereas a change in AFD computed from the highest b-value shell could more directly reflect a change in the underlying *f*, reducing the potentially confounding effect of discrepancies with the response function. The addition of noise had negligible effects on these relationships (Figure S3), however, the estimated AFD appeared to be more variable with decreasing SNR (greater noise).

### 3.2. *In vivo* data

#### 3.2.1. Impact of b-value sampling scheme

In order to assess the impact of b-value sampling-schemes on tract-specific age relationships, we visualise our data as a heatmap (Figure 3; S4). The coefficient of determination (R^2^) derived from the linear model for each tract is organised into hierarchical clusters with branching dendrograms.

**Figure 3:**
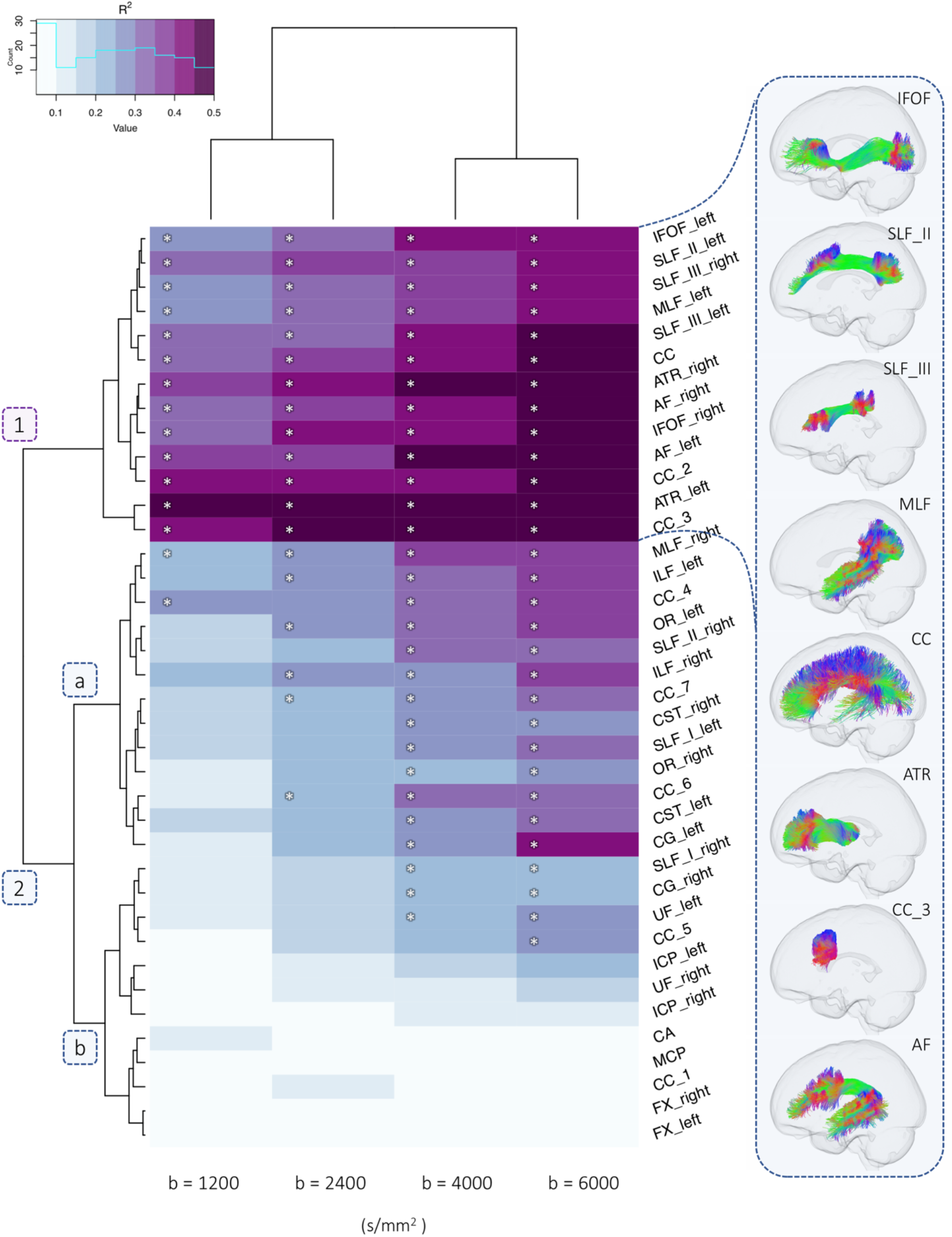
Dendrogram heatmap highlighting clusters of tracts which differentially describe age-related differences in apparent fibre density (AFD) across various single-shell b-value sampling schemes. Heatmap colour intensity reflects range of R^2^ values derived from a linear model including age, sex, and RMS displacement. Significant age-effects (*p_FWE_* < .05) are annotated with an asterisk (*). A depiction of several fibre pathways in one cluster is presented on the right.

##### Single-shell single-tissue FBA

We observe two main clusters of single-shell b-value sampling-schemes: the first including low and moderate diffusion-weightings (b = 1200; 2400 s/mm^2^); and the second including high diffusionweightings (b = 4000; 6000 s/mm^2^). Secondly, two tract-specific hierarchical clusters are observed, represented by branching dendrograms (Figure 3: clusters 1 and 2a,b). Sensitivity to age relationships was improved at high diffusion-weightings (Table 1), whereby AFD exhibited a significantly stronger relationship with age at ss_6000_ compared with: ss_4000_ (6/38 tracts); ss_2400_ (25/38 tracts); and ss_1200_ (27/38 tracts).

**Table 1:** Variance in AFD explained by age for each single-shell sampling scheme across tracts.

| tracts |  | R<sup>2</sup> |  |  |  | R<sup>2</sup> Difference [95% CI] |  |  |  |  |  |
| --- | --- | --- | --- | --- | --- | --- | --- | --- | --- | --- | --- |
|  |  | SS6000 | SS4000 | SS2400 | SS1200 | SS6000 > SS4000 |  | SS6000 > SS2400 |  | SS6000 > SS1200 |  |
| AF | L | .50 | .49 | .39 | .35 | .01 | [-.11, .06] | .12 | [-.01, .18] | **.15** | [.07, .23] |
|  | R | .46 | .43 | .37 | .33 | .02 | [-.08, .09] | .08 | [-.03, .13] | **.13** | [.03, .21] |
| ATR | L | .53 | .50 | .43 | .42 | .03 | [-.15, .11] | .11 | [-.05, .20] | **.11** | [.02, .27] |
|  | R | .48 | .42 | .38 | .36 | .06 | [-.01, .13] | **.10** | [.01, .21] | .12 | [-.05, .21] |
| CA |  | .06 | .03 | .03 | .11 | .04 | [-.01, .12] | .03 | [-.06, .12] | -.05 | [-.17, .13] |
| CC | full | .44 | .39 | .32 | .31 | .04 | [-.07, .10] | **.12** | [.03, .20] | .13 | [-.04, .21] |
|  | 1 | .05 | .02 | .02 | .05 | .03 | [-.04, .13] | .03 | [-.04, .17] | .01 | [-.12, .14] |
|  | 2 | .45 | .43 | .38 | .38 | .03 | [-.13, .10] | .07 | [-.07, .17] | .07 | [-.04, .19] |
|  | 3 | .48 | .46 | .43 | .42 | .02 | [-.03, .07] | **.05** | [.01, .14] | .06 | [-.03, .14] |
|  | 4 | .35 | .31 | .24 | .25 | .04 | [-.02, .08] | **.11** | [.05, .21] | .10 | [-.03, .17] |
|  | 5 | .22 | .16 | .11 | .06 | **.06** | [.02, .10] | **.12** | [.06, .19] | **.16** | [.07, .23] |
|  | 6 | .34 | .29 | .21 | .14 | .04 | [-.02, .08] | **.13** | [.07, .20] | **.19** | [.10, .31] |
|  | 7 | .31 | .29 | .22 | .18 | .02 | [-.04, .08] | **.09** | [.04, .15] | **.13** | [.05, .20] |
| CG | L | .38 | .27 | .18 | .10 | .11 | [-.03, .23] | **.20** | [.06, .34] | **.28** | [.11, .41] |
|  | R | .21 | .20 | .11 | .06 | .01 | [-.14, .10] | .10 | [-.02, .20] | **.16** | [.05, .29] |
| CST | L | .34 | .27 | .19 | .16 | .07 | [-.01, .15] | **.15** | [.06, .21] | **.18** | [.05, .27] |
|  | R | .29 | .28 | .20 | .15 | .01 | [-.08, .09] | .09 | [-.01, .17] | **.14** | [.05, .26] |
| FX | L | .06 | .03 | .01 | .01 | .02 | [-.04, .10] | .04 | [-.02, .12] | .05 | [-.01, .16] |
|  | R | .05 | .02 | .01 | .01 | .03 | [-.04, .10] | .05 | [-.04, .16] | .03 | [-.12, .18] |
| ICP | L | .21 | .18 | .11 | .04 | .03 | [-.01, .14] | **.11** | [.02, .23] | **.17** | [.02, .31] |
|  | R | .11 | .11 | .07 | .08 | -.01 | [-.07, .07] | .03 | [-.07, .14] | .03 | [-.10, .16] |
| IFOF | L | .44 | .40 | .34 | .29 | .03 | [-.02, .13] | **.10** | [.01, .18] | **.15** | [.09, .26] |
|  | R | .46 | .42 | .40 | .33 | .04 | [-.02, .12] | .06 | [-.01, .14] | **.12** | [.02, .22] |
| ILF | L | .39 | .34 | .27 | .22 | .05 | [-.02, .16] | **.13** | [.05, .22] | **.18** | [.10, .27] |
|  | R | .35 | .26 | .24 | .21 | **.09** | [.01, .19] | **.10** | [.04, .24] | **.14** | [.05, .25] |
| MCP |  | .07 | .06 | .05 | .08 | .01 | [-.03, .05] | .02 | [-.07, .10] | -.02 | [-.13, .15] |
| MLF | L | .43 | .39 | .30 | .26 | **.04** | [.01, .09] | **.13** | [.06, .17] | **.17** | [.10, .24] |
|  | R | .39 | .34 | .28 | .20 | .05 | [-.04, .09] | **.11** | [.02, .18] | **.19** | [.07, .25] |
| OR | L | .36 | .30 | .25 | .18 | **.05** | [.01, .13] | **.11** | [.05, .21] | **.17** | [.10, .28] |
|  | R | .28 | .25 | .19 | .13 | .03 | [-.04, .10] | **.09** | [.02, .17] | **.15** | [.05, .30] |
| SLF_III | L | .47 | .44 | .33 | .30 | **.04** | [.01, .08] | **.14** | [.08, .19] | **.18** | [.11, .26] |
|  | R | .41 | .38 | .31 | .28 | .04 | [-.09, .10] | **.10** | [.01, .18] | **.13** | [.01, .24] |
| SLF_II | L | .41 | .40 | .32 | .29 | .01 | [-.04, .06] | **.09** | [.02, .14] | **.12** | [.06, .20] |
|  | R | .31 | .29 | .21 | .17 | .02 | [-.05, .05] | **.10** | [.04, .15] | **.13** | [.07, .22] |
| SLF_I | L | .29 | .24 | .19 | .15 | **.05** | [.01, .11] | **.10** | [.05, .15] | **.14** | [.05, .22] |
|  | R | .22 | .20 | .15 | .08 | .02 | [-.03, .06] | **.07** | [.02, .13] | **.14** | [.06, .24] |
| UF | L | .27 | .23 | .11 | .10 | .04 | [-.05, .16] | **.16** | [.07, .27] | **.16** | [.03, .28] |
|  | R | .17 | .09 | .04 | .04 | .08 | [-.01, .21] | **.13** | [.03, .26] | **.13** | [.02, .32] |

*Note:* R<sup>2</sup> represents the multiple coefficient of determination computed using a linear model for each tract, with age, sex and motion as predictors. Difference indicates difference between R<sup>2</sup> coefficients. Square brackets show 95% bias corrected accelerated (BCa) confidence intervals computed with 10,000 bootstrapped samples. Bold=differences in R<sup>2</sup> where 0 was not captured by the confidence intervals. Abbreviations: AF: arcuate fasciculus; ATR: anterior thalamic radiation; CA: anterior commissure; CC: corpus callosum [1=rostrum, 2=genu, 3=rostral body, 4=anterior midbody, 5=posterior midbody; 6=isthmus, 7=splenium]; CG = cingulum; CST: corticospinal tract; FX: fornix; ICP: inferior cerebellar peduncle; IFOF: inferior fronto-occipital fasciculus; ILF: inferior longitudinal fasciculus; MCP: middle cerebellar peduncle; MLF: middle longitudinal fasciculus; OR: optic radiation; superior longitudinal fasciculus: SLF [I, II, III]; UF: uncinate fasciculus.

The first tract cluster is composed of a sub-cluster of regions where a high proportion of age-related variance is described across all diffusion weightings (median R^2^ = .40). The first sub-cluster (Figure 3: cluster 1) includes several association tracts (left MLF, bilateral IFOF, left SLF II, bilateral SLF III, bilateral ATR, bilateral AF) and commissural tracts (corpus callosum: full extent, genu, rostral body). Significant age-relationships are observed for all of the sampling schemes (b = 1200; 2400; 4000; 6000 s/mm^2^), with an increase in the estimated R^2^ when going to higher diffusion weightings (Figure 4). The proportion of variance explained for the high diffusion-weightings (b = 4000 and 6000 s/mm^2^) ranged from 38% to 53% (Table 2). Despite the consistent sensitivity to age-related development in this tract cluster, a greater b-value dependence on these relationships was observed when moving from high to low b-values, particularly for association tracts such as bilateral SLF III, left SLF I, left IFOF and left MLF.

**Figure 4:**
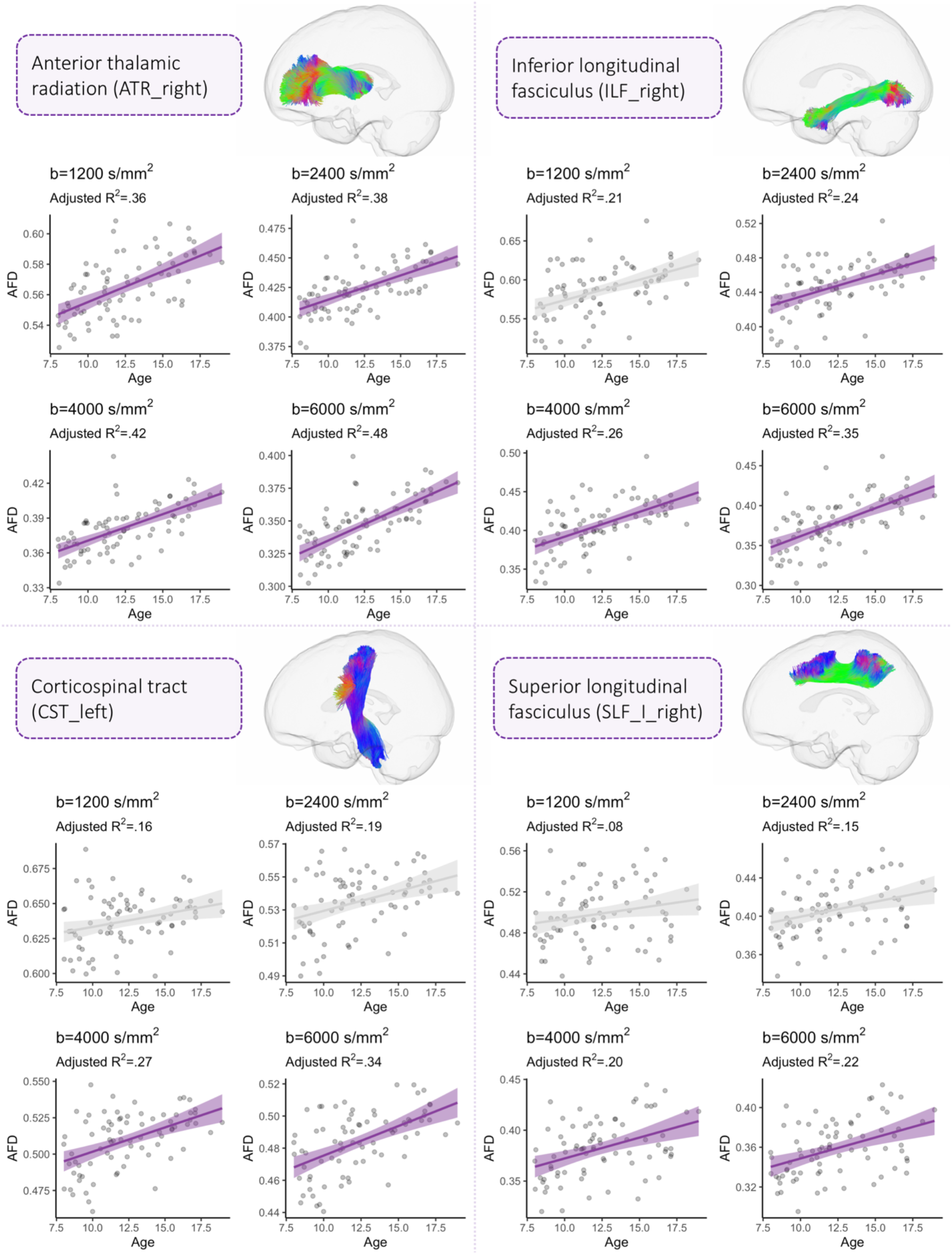
The relationship between AFD and age across four regions including: the right anterior thalamic radiation (ATR_right), inferior longitudinal fasciculus (ILF_right), corticospinal tract (CST_left), and superior longitudinal fasciculus I (SLF_I_right). Each region is representative of individual tract clusters where a progressive increase in the coefficient of determination (R^2^) is observed when moving from low to high diffusion-weightings. Sampling schemes whereby AFD was significantly associated with age are coloured in purple (*p_FWE_* < .05).

The second tract cluster is composed of a sub-cluster (Figure 3: 2a) of association tracts (left SLF_I, right SLF_II, bilateral ILF, right MLF, left CG, bilateral OR), projection tracts (bilateral CST), and commissural tracts (corpus callosum: anterior midbody, isthmus). In this sub-cluster, significant age-relationships are predominantly observed at high diffusion-weightings where the proportion of variance ranged from 24% to 39%, compared with at low-to-moderate diffusion-weightings (10% to 28%). The second sub-cluster (Figure 3: 2b) includes cerebellar tracts (MCP, bilateral ICP), rostrum of the corpus callosum, bilateral fornix, right SLF_I and bilateral UF. This represented a sub-cluster of tracts which captured little-to-no variation across development across moderate-low b-values (1% to 15%).

##### Multi-shell multi-tissue FBA

Consistent with the single-shell single-tissue results, sensitivity to age relationships was improved at high diffusion-weightings (Table S1). We observed two main clusters of multi-shell b-value sampling schemes; the first including multiple combinations of low, moderate, and high b-value sampling schemes, and the second including various combinations of high b-value sampling schemes (Figure S5). In addition, we observed two main tract-clusters consistent with the single-tissue results: the first including various left-lateralised association tracts and corpus callosum projections; and the second including predominantly cerebellar tracts, projection tracts (CST) and association tracts (including right SLF_II, SLF_I, ILF, CG, and OR). Overall, we observed a general reduction in the proportion of detectable age-related variance when adding multiple shells for AFD estimation across various tracts.

#### 3.2.2. Whole brain fixel-based analysis

In order to evaluate the sensitivity of FBA to age-related microstructural development across sampling-schemes, we performed four separate statistical analyses. For each single-shell sampling scheme (b = 1200; 2400; 4000; and 6000 s/mm^2^) we tested the relationship between age and AFD using the CFE method (Raffelt, et al., 2015).

FBA revealed a significantly positive relationship between AFD and age across all b-values (*p_FWE_* < .05). No significant age effects were observed in the opposite direction (*p_FWE_* > .05). We observed a general decrease in the number of significant fixels (n_sig_) when moving from high to low b-values (ss_6000_: n_sig_ = 13,382; ss_4000_: n_sig_ = 10,070; ss_2400_: n_sig_ = 7,283, ss_1200_: n_sig_ = 5,506). In terms of anatomical overlap between results, 58% of significant fixels overlapped between ss_6000_ and ss_4000_, 43% of significant fixels overlapped between ss_6000_ and ss_2400_; and 20% of significant fixels overlapped between ss_6000_ and ss_1200_. Visualisations of significant and overlapping fixels across diffusion-weightings are depicted in Figure 5. The core regions overlapping across all sampling schemes include the body and splenium of the corpus callosum, left IFOF, left ATR, left SLF, and right CST.

**Figure 5:**
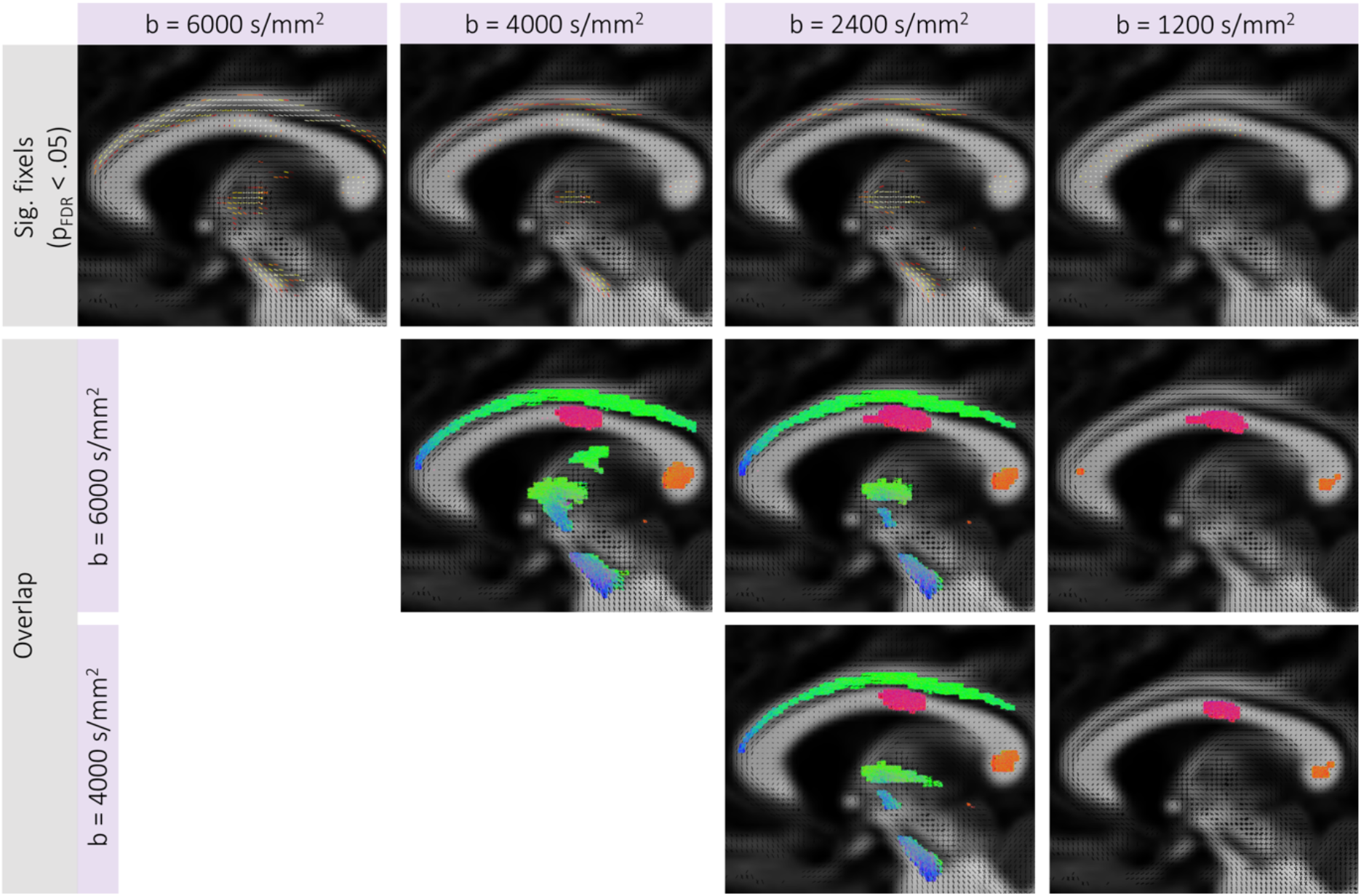
Fixel-based analysis results. Top row displays tracts traversing fixels which are significantly increasing in AFD with age for each b-value sampling scheme (*p_FWE_* < .05). The second (ss_6000_ vs ss_4000_; ss_6000_ vs ss_2400_) and third (ss_4000_ vs ss_2400_) rows display maps of the tracts traversing overlapping fixels between separate FBA results. Results are shown on a representative sagittal slice.

## 4. Discussion

In this study we demonstrate a b-value dependence on estimates of apparent fibre density. Our results highlight that AFD more prominently reflects age-related white matter development at high b-values.

### 4.1. Simulations

The simulations for multiple sampling schemes revealed an improved correspondence between estimated AFD and the underlying intra-axonal fibre properties when using high b-value shells (b = 4000 or b = 6000 s/mm^2^). When moving to lower b-values, or including the complete set of multishell data, we observed a larger dependency of AFD on extra-axonal perpendicular diffusivity. This could suggest that any changes in the true underlying fibre density could be camouflaged by concomitant changes in perpendicular diffusivity, whereby a simultaneous reduction of the intra-axonal volume fraction and *D*_*e*,⊥_ could result in the AFD remaining the same.

AFD is hypothesised to be proportional to the intra-axonal signal fraction of a fibre population (Raffelt, et al., 2012). With increasing b-value, the intra- and extra-axonal signal is differentially attenuated, leading to greater signal contribution from the intra-axonal space (Tournier, et al., 2013). Therefore, an increase in AFD can suggest alterations to axonal properties, such as axon count, packing density, and diameter (Raffelt, et al., 2017). However, our results suggest that AFD is dependent on the extra-axonal signal when including lower b-values, as the mismatch between estimated AFD and simulated intra-axonal signal fraction across varying *D*_*e*,⊥_ is exaggerated.

As such, a change in AFD estimated at high diffusion-weightings (in this case b = 4000 or 6000 s/mm^2^) could more directly reflect a change in the underlying axon density compared with lower b-value shells or multi-shell acquisitions, reducing the potential confounding effect of discrepancies with the response function.

### 4.2. *In vivo* data

When considering *in vivo* developmental data, the dependence of b-value on estimates of AFD was reflected by improved sensitivity to age relationships. Several association tracts consistently described age-related differences in AFD across moderate to high diffusion-weightings, including the left MLF, bilateral IFOF, left SLF II, bilateral SLF III, bilateral ATR, bilateral AF, and anterior segments of the corpus callosum. These regions, particularly the corpus callosum, arcuate and superior longitudinal fasciculus, appear to be sensitive to age-related differences in microstructure regardless of dMRI acquisition scheme or analysis technique (Genc, et al., 2018b; Ladouceur, et al., 2012; Lebel and Beaulieu, 2011; Sawiak, et al., 2018). Our results suggest that this sensitivity to developmental effects is significantly improved with increasing b-value, particularly for the SLF and posterior segments of the corpus callosum.

A group of left-lateralised association tracts (e.g. left CG, MLF, OR, SLF_III, SLF_I, IFOF) better described age-related variance in AFD when comparing the highest b-value (b=6000 s/mm^2^) with high to moderate b-values (b = 4000 or 2400 s/mm^2^). Left-lateralisation of language has been well documented (Catani, et al., 2005) and related to microstructure (Lebel and Beaulieu, 2009). The microstructure of lateralised association tracts is likely linked with the ongoing development of complex cognitive processes throughout childhood and adolescence (Blakemore and Choudhury, 2006; Jung and Haier, 2007). Our results suggest that lateralised association tracts linked with language and cognitive development are better characterised at high b-values. This is likely due to improved sensitivity and specificity to axonal microstructure in the branching endpoints of these tracts integrating such higher order functions across fronto-parietal, fronto-occipital, and occipitotemporal pathways. Future work should focus on investigating subject-specific branching endpoints of these tracts, to assess individual variation in microstructure.

One key observation was that a higher proportion of age-related variance was observed in the single-tissue analyses compared with the multi-tissue analyses. A decrease in discriminative power of age-related development was observed across a number of multi-shell configurations, more heavily weighted towards those which included low-to-moderate b-values. It is possible that singleshell analyses at higher b-values may better isolate the true effect of changing intra-axonal properties, and not clutter it with mismatches of the response function and/or other effects in the extra-axonal space. Future work comparing the current approach with emerging methods such as single-shell three-tissue CSD (Aerts, et al., 2019; Dhollander, et al., 2019) and simultaneous voxelwise estimation of the response function and fibre orientations (Jespersen, et al., 2007) are warranted to explore this further.

The results of the whole-brain FBA revealed a b-value dependence on age-related differences in AFD. Notably, more widespread associations with age were observed at high diffusion-weightings, implicating a number of regions which were not found using other sampling-schemes. This b-value dependence suggests that whilst some core regions such as the body and splenium of the corpus callosum are clearly exhibiting strong age-related development across all sampling schemes, a degree of anatomical sensitivity and specificity is lost at lower diffusion-weightings. This is not to say that studies performing FBA with low-to-moderate b-values will completely lose sensitivity to age-related effects or clinical group differences. However, in conditions with subtle differences in underlying neurobiology or microstructure, going to higher b-values may improve the characterisation of AFD and thus improve the detectability of clinically relevant group differences.

Overall, AFD derived from high b-values (b = 4000 or 6000 s/mm^2^) best modelled age-relationships for the majority of white matter tracts tested. These results, combined with the simulations, suggest that axonal properties (such as axon density) dominate age-related variance in AFD at high b-values, whereas extra-axonal signal contamination at decreasing diffusion-weightings incrementally suppress this effect.

### 4.3. Implications

Our results bear implications for fixel-based analysis applications using retrospectively collected dMRI data which may not be optimal for the estimation of AFD. The biological interpretation of group differences in AFD should be tailored to the acquisition scheme used. Relative differences in AFD at high b-values could relate to the true underlying axon density; at moderate b-values could relate to overall white matter fibre density; and at low b-values could relate to white matter fibre density including potential extra-axonal signal contamination.

Although we have demonstrated a clear b-value dependence on developmental patterns of AFD, it is important to note that the *in vivo* results at high diffusion-weightings are specific to the ultra-strong gradient system used here, resulting in a higher SNR compared with what could be a achieved on a standard MR system. Promisingly, our simulation results suggest that the effect of b-value and discrepancy with the response function dominates the effect of noise (Figure S3), even at a lower SNR which closely matched our *in vivo* data (SNR=50). Therefore, we expect that our observations at high b-values may be reproducible on a standard 3.0T system. As strong gradient systems become increasingly available, the practicalities of acquiring such high quality dMRI data at higher b-values is becoming less cumbersome (Chamberland, et al., 2018; McKinnon and Jensen, 2019; Moss, et al., 2019).

Whilst in this study we have used a developmental population of children and adolescents as an exemplar of a b-value dependence on estimates of AFD, these findings can be applied more broadly and bear implications for a range of group studies (e.g. clinical groups or ageing adults).

### 4.4. Limitations and future directions

One limitation of the current study is that we have no ground truth on the development of axonal density over childhood and adolescence. Therefore, our interpretations of improved intra-axonal signal sensitivity rests on age-relationships investigated here, which has also been used previously (Maximov, et al., 2019; Pines, et al., 2019). Whilst we have attempted to understand how AFD can vary across multiple simulated fibre geometries, we do not know how the underlying fibre properties (such as axon diameter) vary with age. Despite this consideration, a recent study of histological validation suggests that AFD is a reliable marker of axonal density in the presence of axonal degeneration (Rojas-Vitea, et al., 2019). This is a promising indicator of the neurobiological properties proportional to AFD. Future work should adopt multi-dimensional approaches to extract meaningful components (Chamberland, et al., 2019), enhance data quality (Alexander, et al., 2017) and harmonise existing data (Maximov, et al., 2019; Tax, et al., 2019).

## 5. Conclusion

We summarise our findings with three main conclusions: (1) the correspondence between apparent fibre density and simulated intra-axonal signal fraction is improved with high b-value shells; and (2) AFD better reflects age-related differences in axonal microstructure with increasing b-value (b = 4000 or 6000 s/mm^2^) over childhood and adolescence. Together, our results suggest that axonal properties dominate the variance in AFD at high b-values.

## 7. Declaration

All authors disclose no real or potential conflicts of interest.

## 8. Acknowledgements

We are grateful to the participants and their families for their participation in this study. We would like to thank Umesh Rudrapatna, John Evans, Alison Cooper, Peter Hobden, Daniel Burley, and Isobel Ward (CUBRIC) for their support with data acquisition. We also thank Gareth Ball and Charles Malpas for statistical advice.

These data were acquired at the UK National Facility for In Vivo MR Imaging of Human Tissue Microstructure funded by the EPSRC (grant EP/M029778/1), and The Wolfson Foundation. DKJ is supported by a Wellcome Trust Investigator Award (096646/Z/11/Z) and a Wellcome Trust Strategic Award (104943/Z/14/Z). SG was supported by an International Society for Magnetic Resonance in Medicine (ISMRM) research exchange grant. CMWT was supported by a Rubicon grant (680-50-1527) from the Netherlands Organisation for Scientific Research (NWO). ER was supported by a Marshall Sherfield fellowship. MC was supported by the Postdoctoral Fellowships Program from the Natural Sciences and Engineering Research Council of Canada (PDF-502385-2017).

## 9. Supplementary material

### 9.1. Supplementary information

All of the children included in the study were typically developing. None of the children reported a previous clinical diagnosis of ADHD or learning disabilities (based on parent report). Out of the children included, we only collected handedness information for 59/78 participants (76%). Of these children, 11 were left-handed (18%) and 48 were right-handed (62%).

### 9.2. Supplementary figures

**Figure S1:**
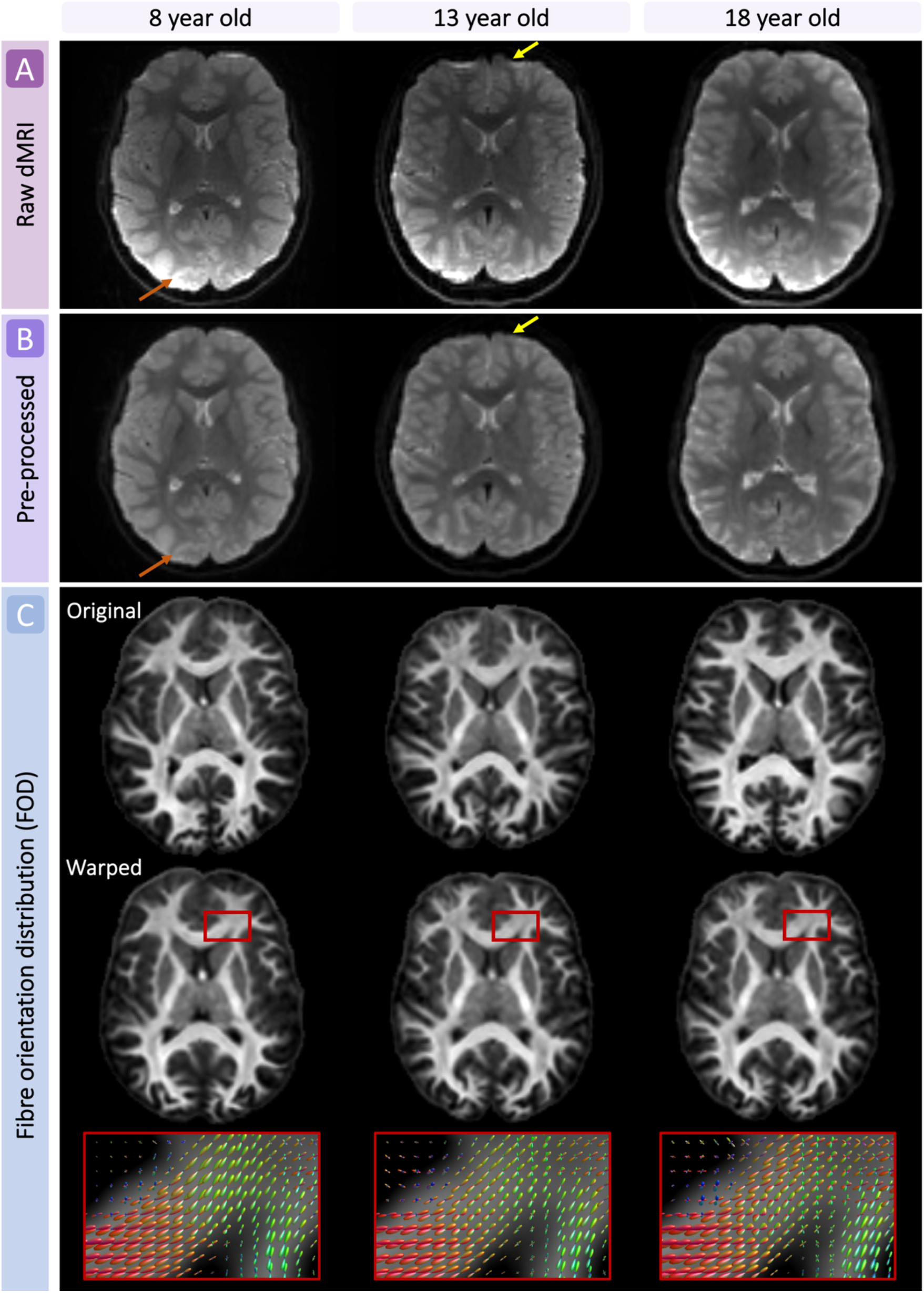
Example of pre-processing and analysis outputs for a representative participant aged 8, 13 and 18 years old. A) Raw dMRI data; B) pre-processed dMRI data; C) fibre orientation distributions (FODs) pre- and post-registration to the population-based template.

**Figure S2:**
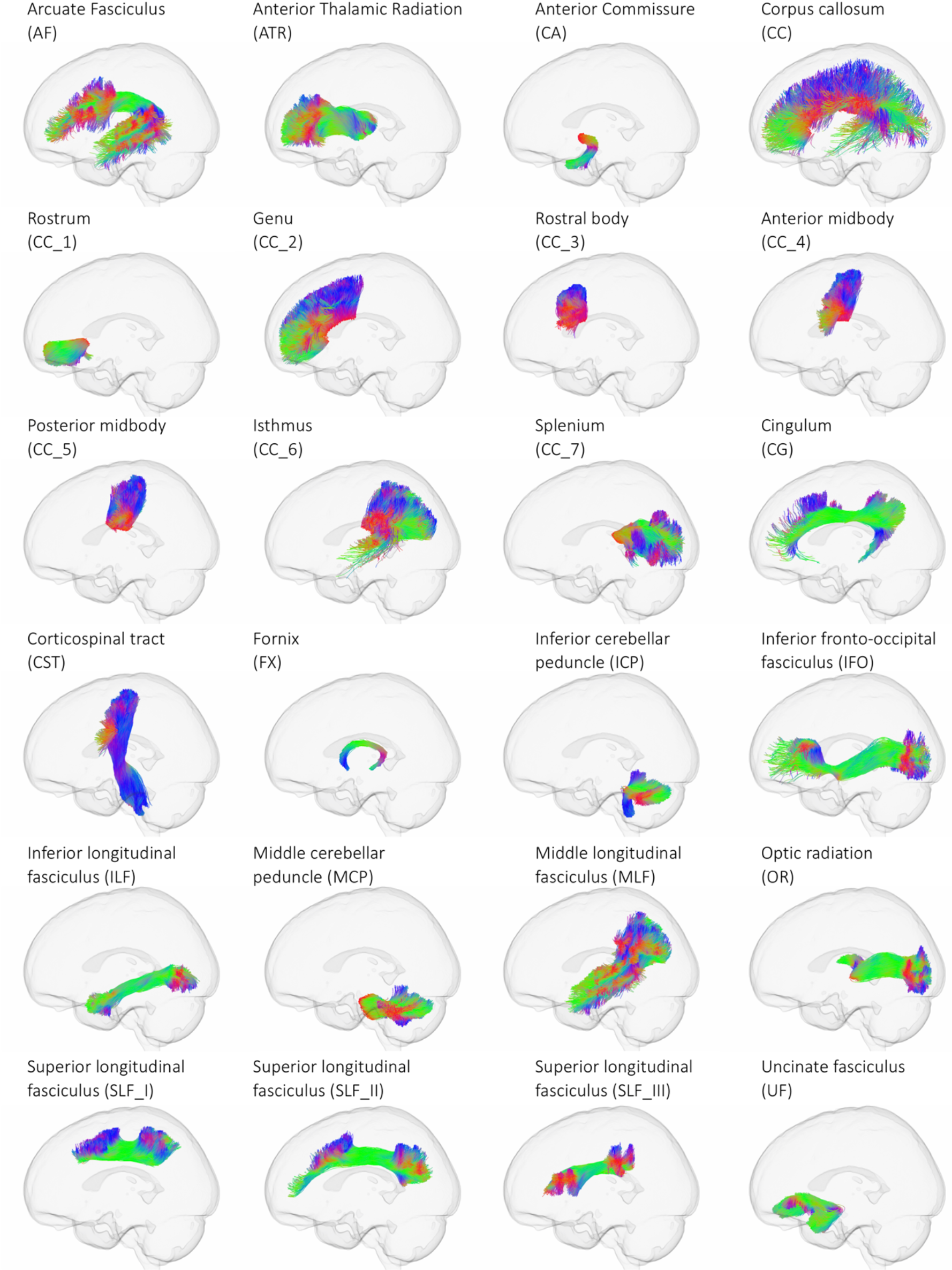
Representative images of tract bundles extracted for statistical analysis, generated using TractSeg. The left view of the tract is presented in each case. Tracts are coloured by the direction of streamlines (red: left-right; green: anterior-posterior; blue: inferior-superior).

**Figure S3:**
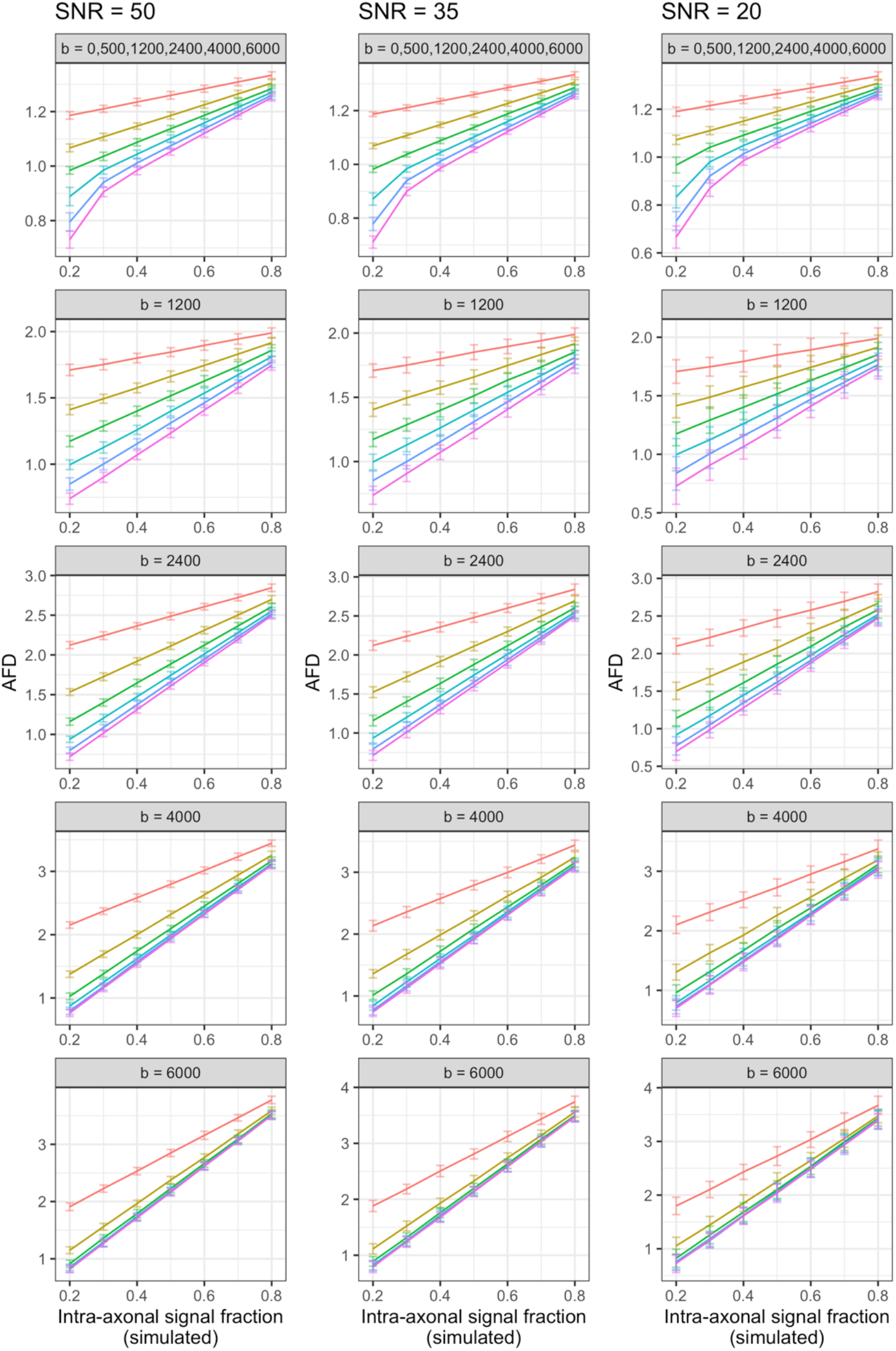
AFD for simulated fibre geometries across five sampling schemes with noise. Variations to simulated intra-axonal signal fraction and perpendicular diffusivity of the extra-axonal space (*D*_*e*,⊥_) were tested to compare AFD across multiple fibre geometries. Simulations were performed with noise (SNR=50; 35; and 25) with 100 Rician noise generalisations (error bars denote mean ±2SD). Sampling schemes reflect the chosen b-values, in s/mm^2^. With lower SNR (greater noise), the estimated AFD was more variable indicated by a larger spread of values, particularly at smaller intra-axonal signal fractions.

**Figure S4:**
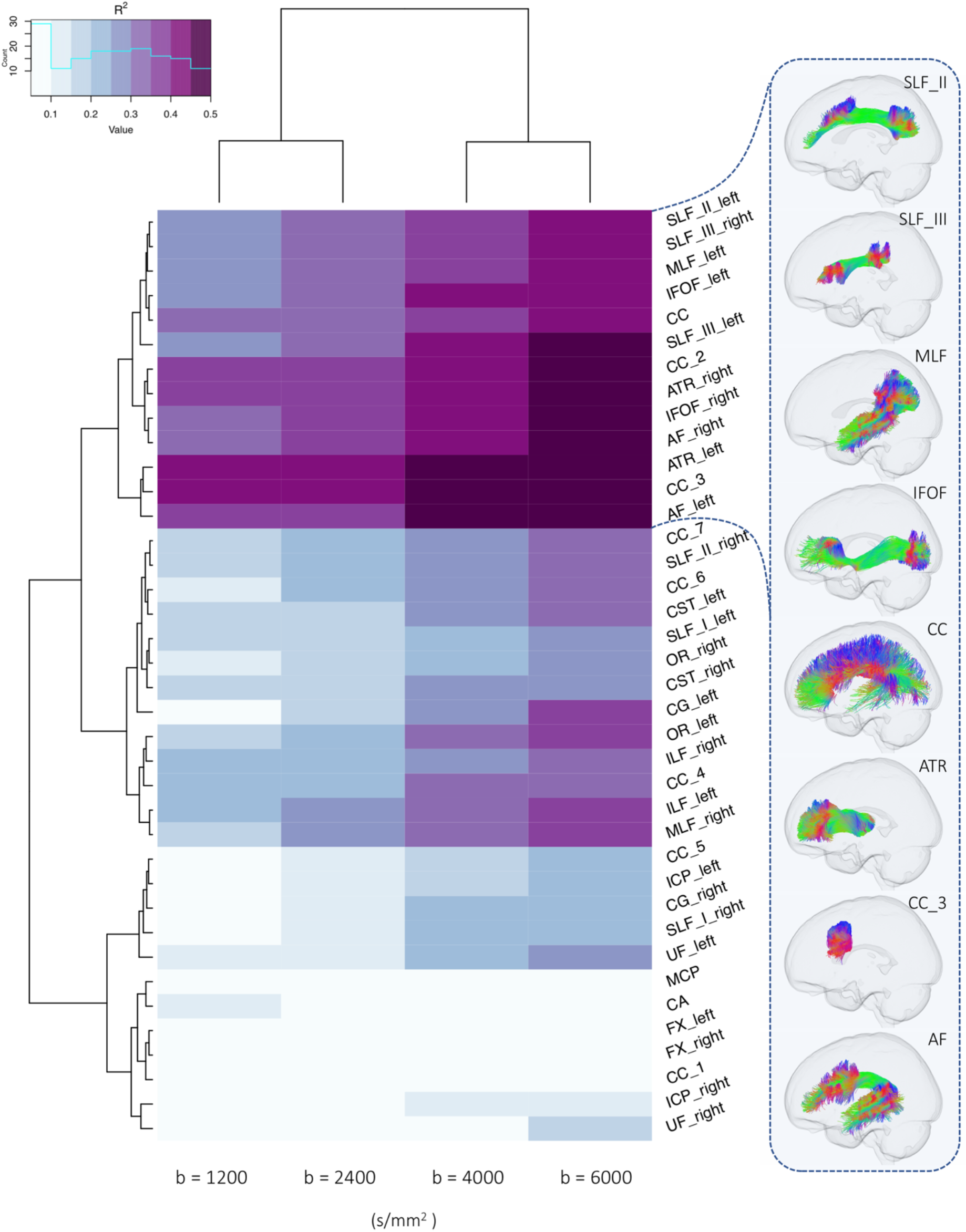
Dendrogram heatmap highlighting clusters of tracts which differentially describe age-related differences in apparent fibre density (AFD) across various single-shell b-value sampling schemes. Heatmap colour intensity reflects range of R^2^ values derived from a linear model including age and sex. Significant age-effects (*p_FWE_* < .05) are annotated with an asterisk (*). A depiction of several fibre pathways in one cluster is presented on the right.

**Figure S5:**
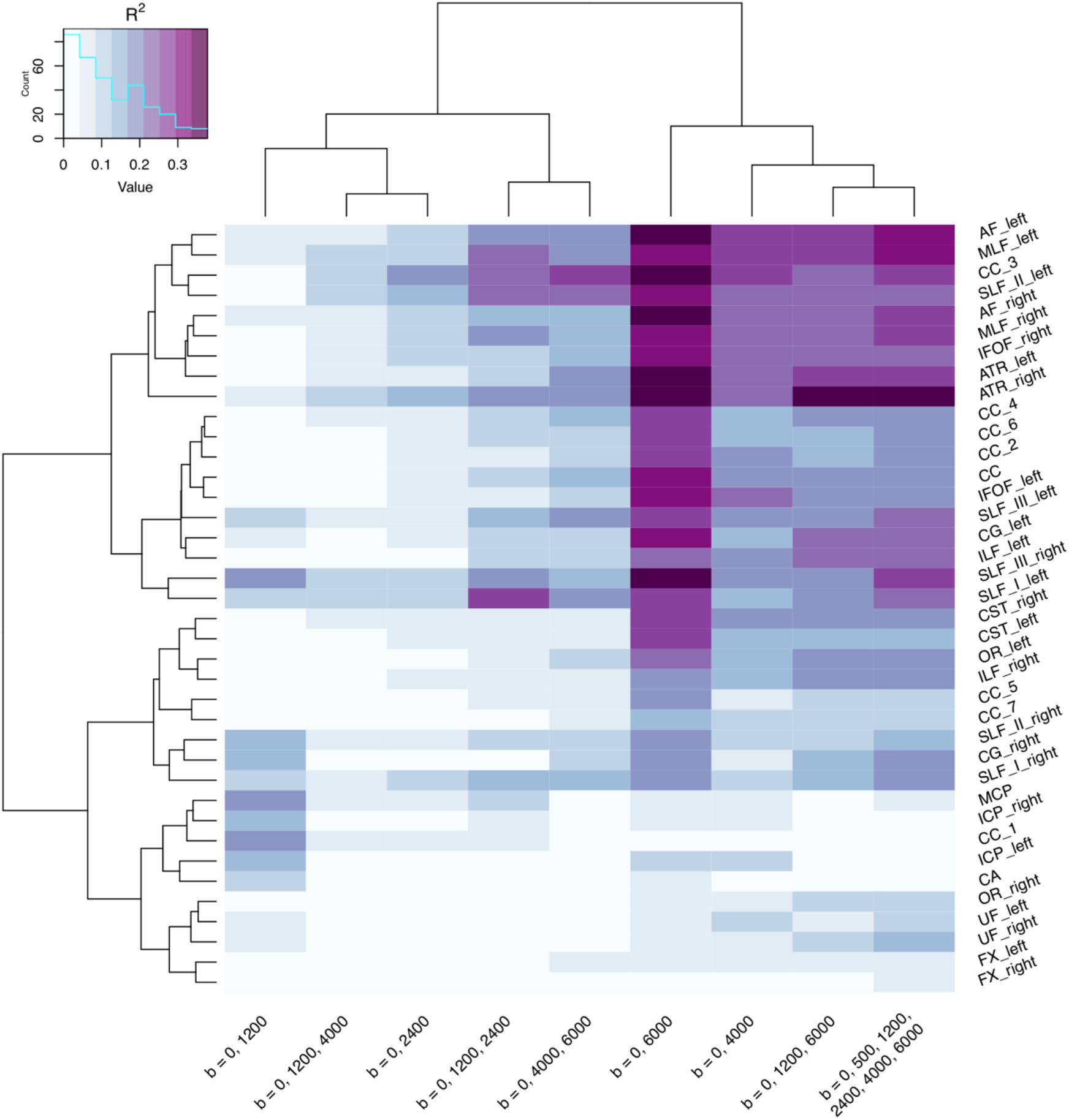
Dendrogram heatmap highlighting clusters of tracts which differentially describe age-related differences in apparent fibre density (AFD) across various multi-shell b-value sampling schemes. Heatmap colour intensity reflects range of R^2^ values derived from a linear model including age and sex. Note: datasets with two diffusion weightings (including the b=0) were processed using the multi-shell multi-tissue FBA framework, thus resulting in separate but comparable results to the single-shell single-tissue FBA (presented in Figure 3).

### 9.3. Supplementary tables

**Table S1:** Variance in AFD explained by age for a set of multi-shell sampling schemes across tracts.

| tracts |  | R<sup>2</sup> |  |  |  |  | R<sup>2</sup> Difference [95% CI] |  |  |  |  |  |
| --- | --- | --- | --- | --- | --- | --- | --- | --- | --- | --- | --- | --- |
|  |  | ms<sub>all</sub> | ms<sub>6000</sub> | ms<sub>4000</sub> | ms<sub>2400</sub> | ms<sub>1200</sub> | ms<sub>6000</sub> > ms<sub>4000</sub> |  | ms<sub>6000</sub> > ms<sub>2400</sub> |  | ms<sub>6000</sub> > ms<sub>1200</sub> |  |
| AF | L | .32 | .35 | .29 | .09 | .08 | .16 | [.01, .16] | **.27** | [.14, .36] | **.27** | [.03, .51] |
|  | R | .29 | .36 | .23 | .11 | .08 | **.21** | [.07, .21] | **.25** | [.14, .38] | .28 | [-.06, .41] |
| ATR | L | .29 | .37 | .24 | .07 | .02 | **.25** | [.07, .25] | **.30** | [.24, .44] | **.36** | [.10, .51] |
|  | R | .39 | .38 | .25 | .14 | .07 | **.24** | [.08, .24] | **.24** | [.13, .34] | **.32** | [.14, .52] |
| CA |  | .02 | .05 | .01 | .02 | .10 | .14 | [-.06, .14] | .04 | [-.08, .12] | -.04 | [-.23, .11] |
| CC | full | .01 | .31 | .19 | .07 | .02 | **.21** | [.08, .21] | **.25** | [.16, .35] | **.30** | [.14, .50] |
|  | 1 | .17 | .01 | .02 | .06 | .19 | .03 | [-.20, .03] | -.06 | [-.15, .05] | -.19 | [-.36, .01] |
|  | 2 | .27 | .27 | .17 | .07 | .03 | **.19** | [.03, .19] | **.21** | [.10, .33] | **.24** | [.09, .45] |
|  | 3 | .17 | .37 | .25 | .17 | .04 | **.19** | [.06, .19] | **.20** | [.12, .28] | **.33** | [.19, .57] |
|  | 4 | .12 | .28 | .17 | .06 | .01 | **.19** | [.06, .19] | **.22** | [.13, .35] | **.27** | [.14, .45] |
|  | 5 | .19 | .18 | .08 | .03 | .02 | **.17** | [.05, .17] | **.15** | [.07, .30] | .17 | [-.08, .36] |
|  | 6 | .11 | .25 | .17 | .06 | .02 | **.15** | [.04, .15] | **.19** | [.09, .29] | **.23** | [.07, .43] |
|  | 7 | .20 | .13 | .12 | .03 | .01 | .15 | [-.04, .15] | **.09** | [.02, .20] | .12 | [-.08, .26] |
| CG | L | .25 | .30 | .14 | .05 | .05 | **.27** | [.08, .27] | **.25** | [.14, .45] | .25 | [-.10, .46] |
|  | R | .18 | .19 | .07 | .02 | .13 | **.21** | [.02, .21] | .17 | [-.01, .33] | .06 | [-.25, .28] |
| CST | L | .14 | .27 | .14 | .05 | .02 | **.20** | [.08, .20] | **.22** | [.10, .36] | .25 | [-.03, .39] |
|  | R | .17 | .28 | .18 | .08 | .02 | **.15** | [.05, .15] | **.19** | [.11, .30] | .26 | [.01, .43] |
| FX | L | .06 | .07 | .05 | .03 | .01 | .08 | [-.03, .08] | .05 | [-.02, .11] | .06 | [-.06, .15] |
|  | R | .04 | .04 | .02 | .01 | .02 | .07 | [-.03, .07] | .03 | [-.03, .11] | .02 | [-.08, .15] |
| ICP | L | .02 | .11 | .11 | .01 | .13 | .11 | [-.12, .11] | .11 | [.01, .33] | -.02 | [-.23, .25] |
|  | R | .03 | .06 | .05 | .03 | .15 | .09 | [-.07, .09] | .03 | [-.08, .18] | -.09 | [-.26, .12] |
| IFOF | L | .20 | .30 | .22 | .05 | .02 | **.16** | [.02, .16] | **.25** | [.14, .35] | **.28** | [.15, .44] |
|  | R | .25 | .32 | .23 | .10 | .02 | **.18** | [.04, .18] | **.23** | [.13, .36] | **.30** | [.08, .57] |
| ILF | L | .23 | .24 | .20 | .04 | .01 | .14 | [-.04, .14] | **.20** | [.10, .35] | **.23** | [.03, .46] |
|  | R | .21 | .18 | .13 | .05 | .01 | .13 | [-.04, .13] | **.13** | [.05, .25] | .18 | [-.01, .37] |
| MCP |  | .05 | .05 | .07 | .05 | .21 | .04 | [-.10, .04] | .00 | [-.14, .13] | -.16 | [-.46, -.03] |
| MLF | L | .31 | .31 | .26 | .11 | .04 | .14 | [.01, .14] | **.20** | [.11, .29] | **.27** | [.13, .46] |
|  | R | .27 | .33 | .22 | .10 | .04 | **.20** | [.06, .20] | **.24** | [.13, .33] | **.30** | [.07, .47] |
| OR | L | .20 | .21 | .14 | .04 | .04 | .17 | [.01, .17] | **.18** | [.02, .26] | .17 | [-.09, .36] |
|  | R | .10 | .07 | .07 | .01 | .02 | .10 | [-.03, .10] | .06 | [-.03, .16] | .05 | [-.06, .27] |
| SLF_III | L | .22 | .28 | .20 | .06 | .09 | **.17** | [.02, .17] | **.22** | [.13, .32] | .19 | [-.07, .44] |
|  | R | .26 | .35 | .20 | .12 | .19 | **.24** | [.10, .24] | **.23** | [.12, .37] | .16 | [-.18, .34] |
| SLF_II | L | .23 | .33 | .23 | .13 | .04 | **.14** | [.06, .14] | **.20** | [.14, .27] | **.29** | [.11, .46] |
|  | R | .13 | .17 | .11 | .06 | .14 | **.14** | [.03, .14] | .11 | [.01, .25] | .03 | [-.19, .25] |
| SLF_I | L | .23 | .29 | .16 | .11 | .09 | **.21** | [.08, .21] | **.18** | [.08, .29] | .21 | [-.04, .41] |
|  | R | .18 | .19 | .12 | .09 | .11 | .15 | [.01, .15] | .11 | [-.01, .22] | .08 | [-.15, .40] |
| UF | L | .09 | .06 | .09 | .01 | .05 | .07 | [-.11, .07] | .06 | [-.08, .19] | .01 | [-.26, .17] |
|  | R | .16 | .06 | .08 | .02 | .06 | .04 | [-.14, .04] | .04 | [-.04, .24] | .01 | [-.19, .24] |

*Note:* R<sup>2</sup> represents the multiple coefficient of determination computed using a linear model for each tract, with age and sex as predictors. Difference indicates difference between R<sup>2</sup> coefficients. Square brackets show 95% bias corrected accelerated (BCa) confidence intervals computed with 10,000 bootstrapped samples. Bold=differences in R<sup>2</sup> where 0 was not captured by the confidence intervals. Abbreviations: AF: arcuate fasciculus; ATR: anterior thalamic radiation; CA: anterior commissure; CC: corpus callosum [1=rostrum, 2=genu, 3=rostral body, 4=anterior midbody, 5=posterior midbody; 6=isthmus, 7=splenium]; CG = cingulum; CST: corticospinal tract; FX: fornix; ICP: inferior cerebellar peduncle; IFOF: inferior fronto-occipital fasciculus; ILF: inferior longitudinal fasciculus; MCP: middle cerebellar peduncle; MLF: middle longitudinal fasciculus; OR: optic radiation; superior longitudinal fasciculus: SLF [I, II, III]; UF: uncinate fasciculus.

